# Sex differences in anxiety and threat avoidance in GAD65 knock-out mice

**DOI:** 10.1101/2023.03.16.532980

**Authors:** Michelle Ulrich, Evangelia Pollali, Gürsel Çalışkan, Oliver Stork, Anne Albrecht

**Author notes:** corresponding author: Anne Albrecht Institute of Anatomy, Otto-von-Guericke-University, Leipziger Str. 44, 39120 Magdeburg, Germany.

## Abstract

Anxiety disorders have been linked to a disbalance of excitation and inhibition in a network of brain structures comprising frontal cortical regions, the amygdala and the hippocampus, among others. Recent imaging studies suggest sex differences in the activation of this anxiety network during the processing of emotional information. Rodent models with genetically altered ϒ-amino butyric acid (GABA) neurotransmission allow studying the neuronal basis of such activation shifts and their relation to anxiety endophenotypes, but to date sex effects have rarely been addressed. Using mice with a null mutation of the GABA synthetizing enzyme glutamate decarboxylase 65 (GAD65-/-), we started to compare anxiety-like behavior and avoidance in male vs. female GAD65-/- mice and their wildtype littermates. In an open field, female GAD65-/- mice displayed increased activity, while male GAD65-/- mice showed an increased adaptation of anxiety-like behavior over time. GAD65-/- mice of both sexes had a higher preference for social interaction partners, which was further heightened in male mice. In male mice higher escape responses were observed during an active avoidance task. Together, female mice showed more stable emotional responses despite GAD65 deficiency. To gain insights into interneuron function in network structures controlling anxiety and threat perception, fast oscillations (10-45 Hz) were measured in *ex vivo* slice preparations of the anterior cingulate cortex (ACC). GAD65-/- mice of both sexes displayed increased gamma power in the ACC and a higher density of PV-positive interneurons, which are crucial for generating such rhythmic activity. In addition, GAD65-/- mice had lower numbers of somatostatin-positive interneurons in the basolateral amygdala and in the dorsal dentate gyrus especially in male mice, two key regions important for anxiety and active avoidance responses. Our data suggest sex differences in the configuration of GABAergic interneurons in a cortico-amygdala-hippocampal network controlling network activity patterns, anxiety and threat avoidance behavior.

**Highlights:**

- Role of GABA in sex-specific anxiety endophenotypes demonstrated in GAD65-/- mice
- Sex- and GAD65-dependent alterations in anxiety, social preference and avoidance
- Enhanced in vitro gamma-beta oscillations in ACC slices of GAD65-/- mice
- Increased parvalbumin(+) interneuron counts in ACC slices of GAD65-/- mice
- Reduced somatostatin(+) interneuron counts in dorsal DG and BLA of male GAD65-/- mice

**Graphical Abstract:** 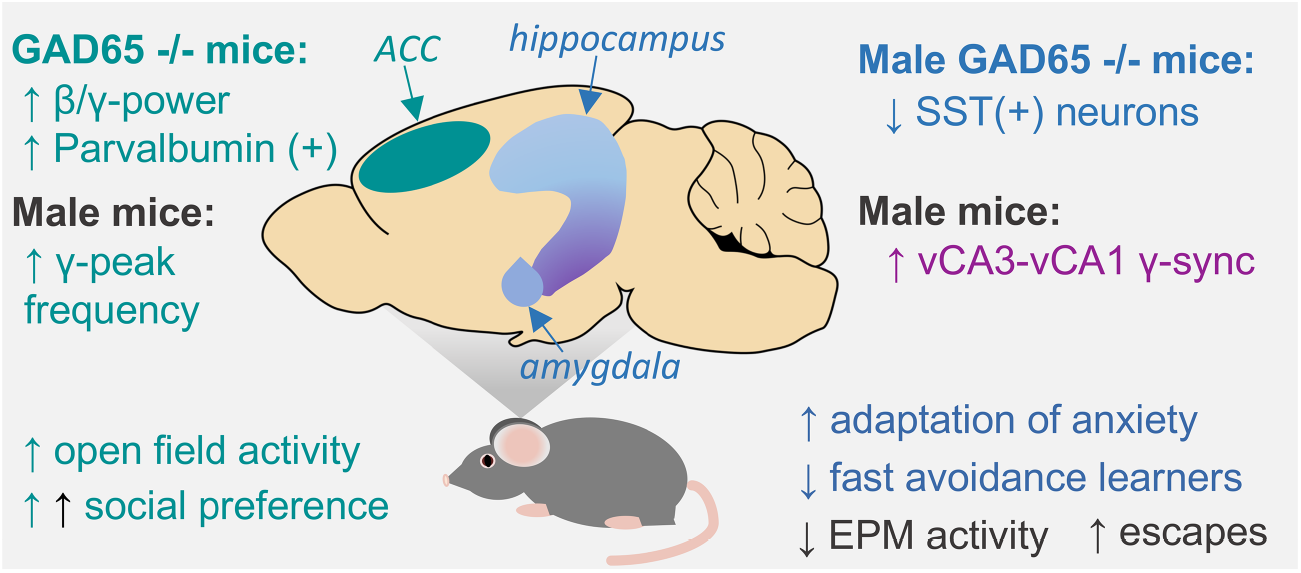

## 1. Introduction

Anxiety disorders are among the psychopathologies with the highest prevalence in western societies and cause enormous individual and socioeconomic burden (Lépine, 2002; Steel et al., 2014). They are characterized by an excessive and uncontrollable worry that interferes with social or occupational functions and is accompanied by sleep disturbances, irritability, restlessness or fatigue (American Psychiatric Association, 2013). Anxiety endophenotypes and cognitive bias to negative interpretations of ambiguous stimuli are strong predisposing factors for developing an anxiety disorder (Ouimet et al, 2009; van der Heiden et al., 2010; Blanco et al., 2014). Further epidemiological studies frequently reveal sex differences (McLean & Anderson, 2009; Vesga-Lopez et al., 2008), indicating a two-times higher prevalence of anxiety disorders in women.

Differential activation of frontal cortical areas such as the medial prefrontal cortex (mPFC) and the anterior cingulate cortex (ACC) is evident between men and women depending on the emotional valence of presented stimuli, induced stress levels or prior adverse childhood experiences (Thomas et al., 2019; Javanbakht et al., 2016; Seo et al., 2017; Stevens & Hamann 2012). Accordingly, a dysfunctional activation of those cortical regions and their interaction with the amygdala and the hippocampal formation are observed during emotion regulation in anxiety patients (Mochkovotich et al., 2014; Goossen et al., 2019; Kim & Kim, 2021). This matches the basic function of these areas and their homologous structures in rodents regulating fear and anxiety as well as emotional affect, attention and decision making (Admon et al., 2013; Tovote et al., 2015; van Heukelum et al., 2020; Chen et al., 2022).

Interactions of cortical frontal areas, amygdala and hippocampus are coordinated through oscillatory rhythms that are under tight control of local inhibitory interneurons (Çalışkan & Stork, 2019). Such GABAergic interneurons control the local excitability and information processing within these regions. Drugs that modulate GABAergic neurotransmission are powerful anxiolytics (Roy-Byrne, 2005) and mice lacking the 65kDa isoform of the GABA-synthetizing enzyme glutamic acid decarboxylase (GAD65) demonstrate a phenotype reminiscent of core symptoms of anxiety disorders, including increased anxiety, hyperarousal, fear generalization and an altered fear extinction (Müller et al., 2014). Different subpopulations of GABAergic interneurons can be described based on their morphological, physiological and neurochemical characteristics. Interneurons containing parvalbumin (PV) are essential for generation of hippocampal oscillations at the gamma frequency range and can modulate anxiety-like behavior and fear persistence (Winkelmann et al., 2014; Çalışkan et al., 2016). Experimental silencing of GABAergic neurotransmission in another subpopulation, somatostatin (SST)-positive interneurons, increases anxiety-like behavior (Miyata et al., 2019; Fee et al., 2021), suggesting a strong contribution of such inhibitory neurons to anxiety endophenotypes.

GAD65-/- mice have been established as model for studying the neurobiological basis of anxiety phenotypes (Kash et al., 1999; Stork et al., 2000), but sex effects have not been yet addressed in these animals. In a previous study we were able to relate a polymorphism in the GAD65 gene to an increased GABA/Glutamate ratios in the ACC, associated with increased anxiety endophenotype and harm avoidance in female but not male participants (Colic et al., 2018). In the current study, we therefore set out to investigate potential sex differences in anxiety-related behavioral tasks and avoidance behavior in a two-way shuttle box in GAD65-/- mice. In addition to the characterization of anxiety-related endophenotypes we investigated beta/ slow gamma oscillations in the ACC as a function of interneuron activity and examined the configuration of GABAergic networks by immunohistochemical labelling of PV- and SST-positive interneurons in frontal cortical, hippocampal and amygdala subregions.

## 2. Material and Methods

### 2.1 Animals

Male and female homozygous GAD65 knock-out (-/-) mice and their wild type (+/+) littermates were obtained from a heterozygous breeding on a C57Bl6/BomTac genetic background, backcrossed for >12 generations, and genotyped by allele-specific polymerase chain reaction (PCR) as described previously (Stork et al., 2000). All mice were bred and raised in the animal facility at the Institute of Biology, Otto-von-Guericke University Magdeburg, and kept in groups of 2–6 of the same sex on an inverse 12 h light/dark cycle (lights on at 7 pm with a 30 min dawn phase) with food and water *ad libitum*. Animal housing and animal experiments were in accordance with European regulations for animal experiments and approved by the Landesverwaltungsamt Saxony-Anhalt (Permission Nr. 42502-2-1177 and -1516). All experiments were performed with young adult female and male mice (age 10-16 weeks) during the dark, active circadian phase with handling under red light illumination by the same female experimenter for each of the three described experiments.

### 2.2 Experiment 1: Behavioral test battery

Female GAD65-/- mice (n=12) and female GAD65+/+ littermates (n=12) as well as male GAD65-/- mice (n=13) and their male GAD65+/+ littermates (n=12) underwent a behavioral test battery consisting of open field (OF) on day 1, elevated plus maze (EPM) on day 2, novel object recognition (NOR) on day 3, social interaction (SI) on day 4 and five days of active avoidance training (AA) on day 7-11. The order of the behavioral tests was chosen according to rising stress levels induced by the single tests.

#### Open field

Mice were placed in the center (25 x 25 cm) of a square arena (50 x 50 cm, bright light illumination, 200 lux) and allowed to explore freely for 20 min. Total distance and distance in center were recorded using the ANY-maze™ Video Tracking System (version 4.50, Stoelting Co., Wood Dale, USA).

#### Elevated plus maze

Mice were placed for 5 min in maze consisting of two closed arms with 15cm high plastic walls and two orthogonally positioned open arms without walls (arm length 35 cm, width 5cm) elevated 110 cm above the floor under low-light conditions (10 lux). Total distance and distance on open arms were recorded using the ANY-maze™ Video Tracking System (version 4.50, Stoelting Co., Wood Dale, USA).

#### Novel object recognition

Following EPM, mice were familiarized to a Y-shaped maze consisting of 3 arms with walls (length 35 cm, width 6cm, height 14 cm) arranged at an 120° angle for 10 min. The next day, two wooden toy blocks were placed at the end of two arms and the test animal was placed in the empty arm to start an 8 min free explorations session under low light conditions (10 lux). The third arm stayed empty and served as starting point for the 8 min exploration. After a 3 h interval in the home cage, one of the wooden toy blocks was randomly interchanged with a Lego plastic ensemble serving as a novel object. All sessions were recorded via ANY-maze™ Video Tracking System (version 4.50, Stoelting Co., Wood Dale, USA) and contact times to objects were scored manually. A discrimination index was calculated as [(contact time novel object/total object contact time) - (contact time familiar object/total object contact time)], with an index close to 1 reflecting strong preference for a novel object, 0 no preference and close to −1 avoidance of a novel object.

#### Social interaction

Social preference and memory were tested in a three-compartment-chamber (chamber: width 40 cm x length 20 cm x height 40 cm) with grid tubes (diameter 8 cm x height 15 cm) in each outer chamber. Mice were first habituated to the arena by allowing free exploration for 5 min, then they were briefly restricted to the central compartment and a same sex interaction partner was introduced to one of the grid tubes. After 5 min of free exploration another new interaction partner was introduced to the second tube to test for social memory in 1 min session. Animals were video tracked using ANY-maze™ software and contact times to tube were scored manually offline. Discrimination indices were calculated for social preference [(contact time mouse tube/ total tube contact time) - (contact time empty tube/ total tube contact time)] and for social memory [(contact time new mouse tube/ total tube contact time) - (contact time familiar mouse tube/ total tube contact time)].

#### Active Avoidance

Animals were placed in a sound isolation cubicle containing a 16x32x20cm acrylic glass arena with a grid floor split in two compartments joined by a door, loudspeaker and ventilation fan (background noise 70dB SPL, light intensity <10 lux; TSE, Bad Homburg, Germany) and received 5 training days. Each training started with a 60 s habituation session to the box, followed by 50 trials consisting of a tone (85 dB, 10 kHz, max. 10 s) followed by footshock (0.4 mA) and a variable intertrial interval of 20 – 40s. Movement of animals was recognized by light beams and shuttling to another compartment during the tone prevented the footshock and was counted as successful avoidance, while shuttling after onset of the footshock was registered as an escape response. Failures, i.e. absence of any shuttling during a trial, was only occasionally observed (<1% of all trials) and not further analyzed. Number of defecation boli as a measure of arousal were counted after each trial.

On day 15, brains were prepared for a ***post-training gene expression analysis*** of somatostatin and parvalbumin in the dorsal ACC, dDG and BLA via qPCR. See Suppl. Material, Fig. S2 for details.

### 2.3. Experiment 2: Oscillations

In a different set of naïve animals of both sexes, slices that contain the anterior cingulate cortex (ACC) were prepared (female GAD65+/+ animals N=8/ slices n=47; female GAD65-/- animals N=6/ slices n=36; male GAD65+/+ animals N=6/ slices n=30; male – GAD65-/- animals N=6/ slices n=30). Mice were decapitated under isoflurane anesthesia and their brains were quickly placed in ice-cold (−4 °C) standard artificial cerebrospinal fluid (aCSF; in mM: NaCl, 129; NaH2PO4, 1.25; glucose, 10; MgCl2, 1; CaCl2, 2; KCl, 3; NaHCO3, 21; bubbled with 95% O2 / 5% CO2, pH≈7.4). Using a vibratome (Campden Instruments), coronal slices of 400 µm were obtained between 1.42 mm and −0.22 mm from Bregma (Paxinos and Franklin, 2001) and transferred to an interface chamber for 1 h incubation at 32 °C perfused with standard aCSF. Then, a modified aCSF containing 50 µM carbachol (CCh), 0.8 µM kainate (KA) and increased K^+^ (∼5 mM) concentration was perfused to induce fast local field potential (LFP) oscillations in the range of beta/gamma band, as described previously (Steullet et al., 2014). Forty-five min later borosilicate glass electrodes of ∼1 MΩ resistance, filled with ACSF, were placed at the dorsal ACC (dACC) of layer 5 and the induced oscillatory activity was recorded for at least 10 min.

The signal was low-pass filtered at 3 kHz with the use of an extracellular amplifier (EXT-02F - Extracellular Amplifier; npi Electronic GmbH, Tamm, Germany). Data were recorded using the Spike2 software with a sampling rate of 5 kHz (CED-Micro1401; Cambridge Electronic Design, Cambridge, UK) and stored on a hard disc for offline analysis.

Artifact-free data from the last two minutes of the recordings were analyzed via computing fast Fourier transformation with a frequency resolution of 2.441 Hz revealing peaks in two frequency ranges at 10-25 Hz (beta oscillations) and 26-45 Hz (slow gamma oscillations). Two oscillation parameters were analyzed: 1) Peak Frequency (Hz): The frequency with the highest power at the corresponding frequency range (10-25 Hz and 26-45 Hz), 2) Integrated Power (mV^2^/Hz): Sum of the binned power values at the corresponding frequency ranges (10-25 Hz and 26-45 Hz). For statistical comparison, only slices showing signals with peak powers higher than 5 µV^2^ and peak frequencies higher than 10 Hz were included.

In an additional set of animals, gamma oscillations were induced in ventral-to-intermediate hippocampal-entorhinal slices via bath-application of 5 µM CCh and recorded in the CA3 and CA1 subregions (see Suppl. Material, Fig. S2 for methodological details).

### 2.4. Experiment 3: Interneuron density

Behaviorally naïve female and male GAD65-/- mice as well as their +/+ littermates of both sexes (n=6 per group) were transcardially perfused with phosphate buffered 4% paraformaldehyde solution (PFA). After overnight incubation in the same fixative and immersion in 30% sucrose solution at 4°C brains were frozen in liquid nitrogen-cooled methyl butane and stored at −20°C. Brains were then cryo sliced into 30 μm serial coronar sections at the level of PFC (1.94-1.54 mm from Bregma), posterior ACC (1.10 - −0.22 mm from Bregma) dorsal hippocampus / amygdala until ventral hippocampus (−1.46 - −2.70 mm from Bregma; Paxinos and Franklin, 2001). Free-floating sections were either incubated with a rabbit monoclonal anti-SST antibody (1:250, Santa Cruz Biotechnology #sc13099, Dallas, Texas, USA) or a combination of a rabbit monoclonal anti-PV antibody (1:500, abcam ab11427, Amsterdam, Netherlands) and a mouse monoclonal anti-GAD67 antibody (1:250; Millipore (Merck KgaA) MAB5406, Darmstadt, Germany) over night. A secondary antibody against rabbit (Alexa-Fluor 488, donkey anti rabbit IgG, 1:1000, Invitrogen (Thermofisher) A21206, Waltham, Massachusetts, USA) was used to label SST and PV, respectively and a biotinylated goat anti-mouse antibody (1:200, Jackson ImmunoResearch, Ely, UK), linked to Cy3 via Streptavidin incubation (1:1000, Jackson ImmunoResearch, Ely, UK), was used to label GAD67. Nuclei were visualized with DAPI (300 nM, Thermofischer, Waltham, MA, USA). Images of the immunostainings were acquired with a DMI6000 epifluorescence microscope (Leica, Wetzlar, Germany) and SST-, PV- and GAD67-positive cells were counted manually within selected brain areas using the software ImageJ. The cell number was normalized to area size of each region of interest (measured with imageJ). The cell counts per target area were averaged from 2 slices per animal covering both hemispheres and the averaged values per individual were used for statistical comparison between groups.

### 2.5. Statistics

Statistical analysis was performed using SPSS 22.0 (IBM Germany GmbH, Ehningen, Germany) and GraphPad Prism (version 9.1.2; GraphPad Software, CA, USA). Effects of sex or genotype and their interaction were analyzed by a two-way analysis of variance (ANOVA) in normally distributed data sets (tested by Shapiro–Wilk test), while in non-normal distributed samples a Mann–Whitney U-test was performed for the main factors sex and genotype. In case of significant interactions, a paired comparison of subgroups was performed using Student’s T-test or Mann-Whitney-U-test. For data sets dependent measures of time and training trials (e.g. Open Field, Active Avoidance), repeated measure ANOVAs (RMA) were performed with sex and genotype as between-subject factors. Here, in case of sphericity issues Greenhouse-Geisser (Epsilon<0.8 at Mauchly’s test of sphericity) or Huynh-Feldt corrections (Epsilon>0.8) were used. Correlation of avoidances on the 2^nd^ training day and fecal boli counts at the last training days were determined using Pearson’s correlation coefficient..

## 3. Results

### 3.1 Sex-specific effects of GAD65 deficiency on anxiety, social preference and active avoidance

Mice first performed an open field test under bright light conditions. The 20 min test phase was analyzed in 5 min bins via repeated measure ANOVA, revealing a significant interaction of time bin and genotype for total distance covered (Fig. 1A; RMA with Huynh-Feldt correction: F(2.633,118.494)=8.783, p<0.001), without additional effects of sex or interactions of sex and genotype. A paired comparisons for genotype in each sex revealed that the main effect of genotype over time was driven by females, with an increased activity at the end of the test the total (Fig. 1B; Student’s T-test for female GAD65-/- vs. GAD65+/+ mice T(22)=3.31, p=0.0049; for male GAD65-/- vs. GAD65+/+ mice: T(23)=0.913, p=0.374). Analyses of the distance covered only in the center revealed interactions of time bin x genotype (Fig. 1C; RMA: F(3,135)=6.22, p=0.001) and of time bin x sex x genotype (F(3,135)=4.026, p=0.009). Paired comparisons for the separate time bins (Fig. 1D) revealed then that especially male GAD65-/- vs. GAD65+/+ mice showed differences in the time course of anxiety-related behavior. While the relative distance in center was increased in male GAD65+/+ compared to male GAD65-/- mice at the beginning of the session (T(23)=2.472, p=0.021), male GAD65-/- mice subsequently increased their exploration of the center area and covered significantly more distance in the center towards the end of the test session (T(23)=-2.785, p=0.011 compared to male GAD65+/+. This differential response in males over the time course of the test drives also the significant main effect of sex observed regardless of time bins (F(1,45) = 4.928, p=0.032). Female GAD65-/- and GAD65+/+ mice displayed comparable levels of activity in the during the first and last 5 min of the open field session.

During a 5 min session on the elevated plus maze under low light conditions on the next day, no genotype effects were observed. Male mice were generally less active than their female counterparts (Fig. 1E, total distance, two-way-ANOVA for effect of sex: F(1,45)=7.65, p=0.008) and no significant effects of genotype or any interactions of sex x genotype were observed. No significant differences in the distance travelled in the open arms were detected (Fig. 1F).

**Figure 1:**
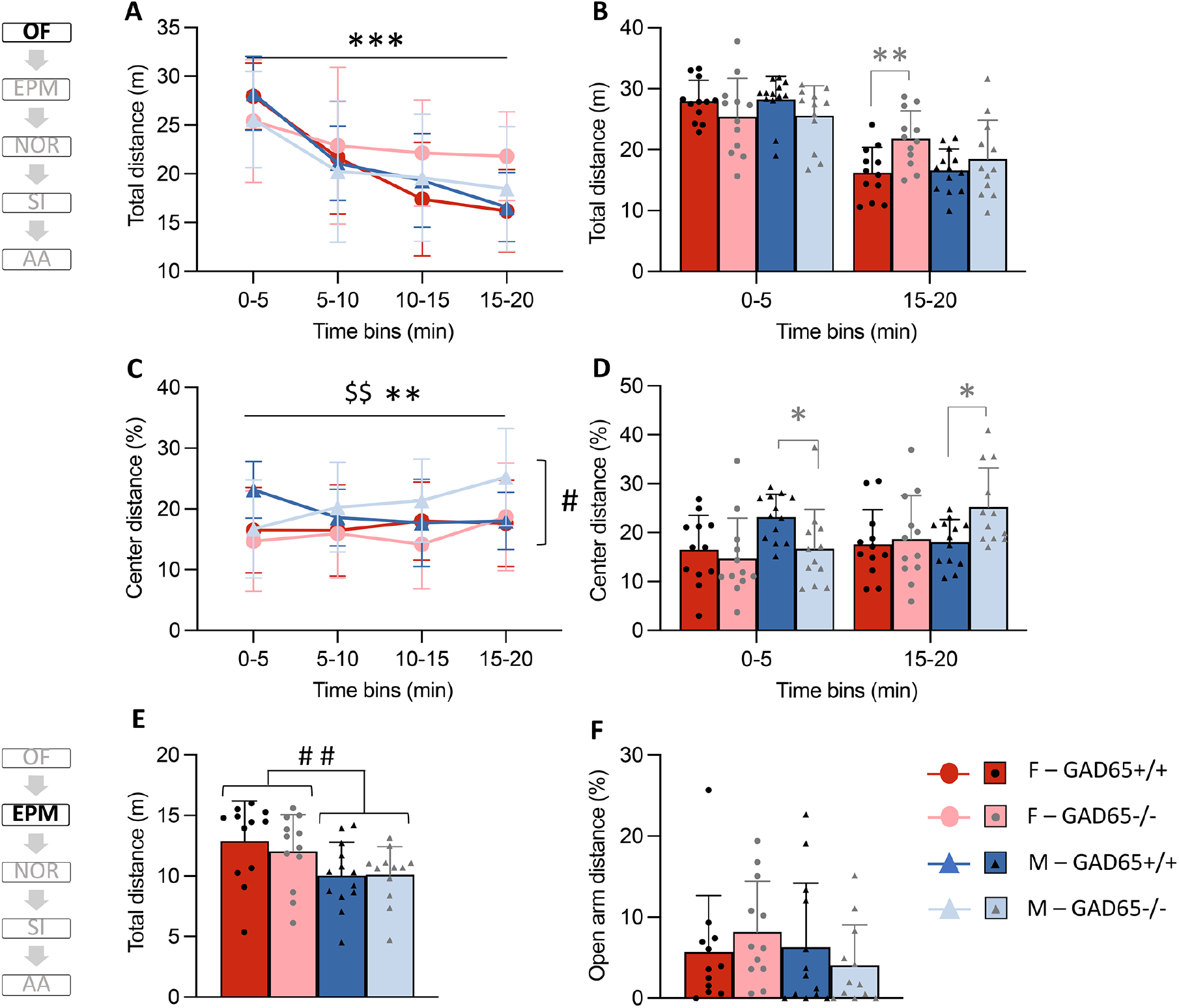
Sex-specific effects of GAD65 deficiency on anxiety. **(A**) In a brightly lit open field, GAD65-/- mice showed increased activity over time compared to their wildtype littermates. **(B)** A detailed analysis of the first and last 5 min of the test revealed that total activity was highest in female GAD65-/- mice towards the end of the test. **(C)** For the activity in the center region, interactions of genotype and sex were observed in addition to an effect of genotype. **(D)** The detailed analysis of the first and last 5 min of the test revealed that male GAD65+/+ mice showed higher activity at the beginning of the open field session, while male GAD65-/- increased their center activity towards the end of the session. Over all time bins, female GAD65+/+ and GAD65-/- mice appear to explore the center less than their male counterparts. **(E)** During an elevated plus maze test under low light conditions, male mice of both genotypes were less active than their female counterparts. **(F)** No significant differences between genotypes and sexes were observed for the activity in the open arms. All data presented as mean ± SD and mean + SD with superimposed individual values for specific analysis time points. Repeated Measure ANOVA (A+C): ** significant interaction of time bin x genotype, p<0.01, *** p<0.001; $$ significant interaction of time bin x sex x genotype, p<0.01; # significant effect of sex, p<0.05; Paired comparison (T-Test; B+D): * (in grey) significant effect of genotype within the male subgroup, p<0.05, ** p<0,01; Two-way ANOVA (E+F): ## significant effect of sex, p<0.01.

Mice were then subjected to a novel object recognition task under low light conditions. Here, no significant effects of genotype or sex or their interactions were observed (Fig. 2A). Since no preference towards a novel object was established in wildtype mice, conclusions regarding recognition memory of male and female GAD65-/- mice are hampered. Discrimination within a social interaction task was achieved well by mice of all groups, with male mice showing higher social preference than females (mouse vs. empty tube; Fig. 2B; two-way-ANOVA for effect of sex: F(1,45)=8.928, p = 0.005). Additionally, GAD65-/- mice showed a greater social preference than their wildtype littermates (two-way-ANOVA for effect of gene: F(1,45)=4.254, p=0.045). Concordantly, the highest social preference was expressed by male GAD65-/- mice. No significant effects were observed for social memory, when preference for an unfamiliar second interaction partner was tested (Fig. 2B).

**Figure 2:**
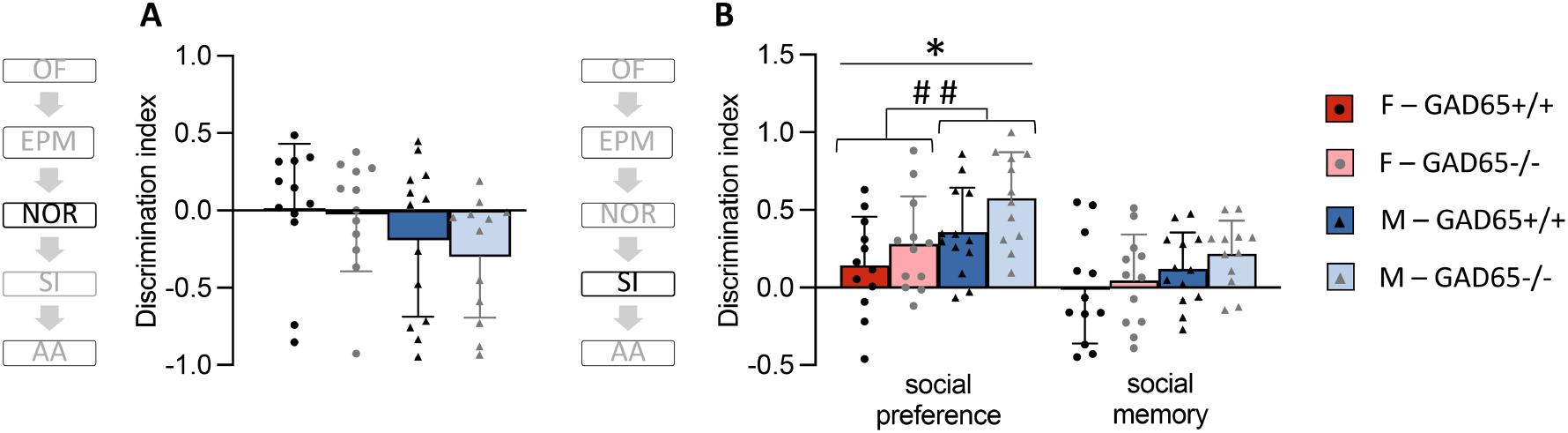
Sex-specific effects of GAD65 deficiency on social preference. **(A)** In the novel object recognition task (NOR) mice explored two similar wooden toy blocks within a Y-maze. After a 3h interval, one of the objects was replaced with a novel object assembled from Lego. Regardless of sex and genotype, all mice did not prefer the novel object. **(B)** However, when mice had to discriminate between test tubes containing an unfamiliar mouse vs. an empty grid tube in the three-chamber social interaction task, all mice spent more time with the social interaction partner. Such social preference was increased in male compared to female mice and GAD65-/- mice compared to their GAD65+/+ littermates regardless of sex. Social memory tested by discriminating between a familiar and unfamiliar social interaction partner was neither affected by sex nor by genotype nor their interaction. All data presented mean + SD with superimposed individual values Two-way ANOVA: * significant effect of genotype, p<0.05; ## significant effect of sex, p < 0.01.

A significant effect of sex was evident during avoidance learning in the shuttle box. Female mice had lower numbers of escapes (Fig. 3A; RMA for escapes with Huynh-Feldt correction with Huynh-Feldt correction: main effect of sex: F(1,43)=4.066, p=0.05; training day x sex: F(3.767,161.978)=1.678, p=0.159). No significant differences were found for the number of successful avoidances over the training days (Fig. 3B). On average, all groups except the male GAD65-/- mice reached more than 50% of successful avoidances on training day 2. Based on previous work in rats investigating the heterogeneity of learning dynamics in a two way-shuttle box (Galatzer-Levy et al., 2014), the 2^nd^ training day was then used to assess such variability within the groups. Here, an animal was classified as a fast learner when it completed more than 50% of successful avoidances (i.e. above chance level) on the 2^nd^ training day. The highest proportion of fast learners was observed in female GAD65+/+ mice (9 out of 12, see Fig. 3C) and the lowest proportion of this subgroup in male GAD-/- mice (3 out of 12). While cognitive control can be established within a few sessions by reaching asymptotic levels of avoidance responses, evidence shows that emotional control and arousal may require longer training (Hadad-Ophir et al., 2017). As an approximation for arousal, defecation boli were counted after each testing session. Significant interactions of training day and sex (Fig. 3D; RMA: F(4,168)=2.58, p=0.039) and of training day x sex x genotype (F(4,168)=2.468, p=0.047), but not of genotype alone were observed. On day 5, paired comparison revealed significantly higher boli counts in male compared to females wildtype mice (Fig. 3E; Student’s T-test: T(23=--2.179, p=0.04), but the number of boli on day 5 did not correlate with the avoidance performance on day 5 (Fig. 3E), indicating that individual differences in avoidance learning did not predict arousal after longer training.

**Figure 3:**
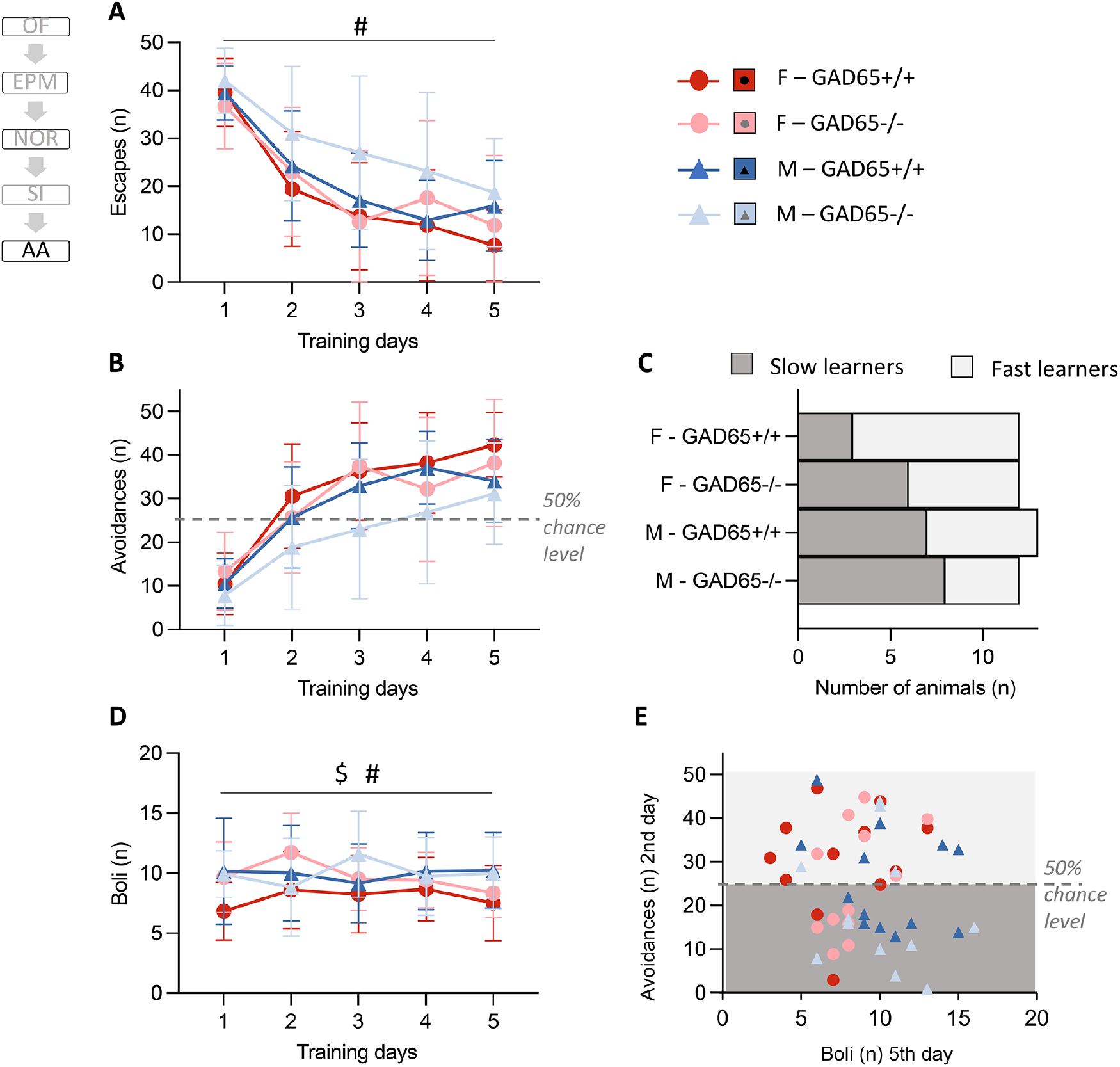
Sex-specific effects of GAD65 deficiency on active avoidance in a 2-way shuttle box. **(A)** Female mice reduced the number of escapes (shuttling after footshock) over five days of training days faster than males. **(B)** The number of avoidances (successful shuttling during tone presentation before footshock onset) was not significantly altered between groups. **(C)** The 2^nd^ day of the training was selected to assess individual variability in avoidance learning by classifying animals into fast learners (> 50% of successful avoidances in the 2^nd^ session) vs. slow learners. The number of fast vs. slow learners are unequally distributed between groups, with male GAD65-/- mice showing only 25% of fast learners (histogram). **(D)** The number of defecation boli after each session as a measure of arousal varied as function of the training days, resulting in a significant effect of genotype and an interaction of sex x genotype. **(E)** On day 5, male GAD65+/+ mice displayed higher boli counts than female wildtype mice. No correlation was observed between boli counts and avoidance performance on the 2^nd^ training day (light grey background: fast learners; dark grey background: slow learners). Data presented as mean ± SD for Repeated Measure ANOVA (A, B, D): # significant effect of sex, p < 0.05; $ significant interaction of sex x genotype, p<0.05.

Taken together, sex differences were evident such as a reduced activity in a low lit elevated plus maze in males. In the active avoidance paradigm, male mice showed more escapes over time and higher boli counts until the end of the training. A mild modulation of sex differences by GAD deficiency was observed in a brightly lit open field. Here, female GAD65+/+ stayed more active towards the end of the test, while male GAD65-/- mice showed a protracted reduction of anxiety-like behavior. They constituted the lowest portion of faster learners in the active avoidance task as well. Female GAD65-/- showed no modulation of anxiety like-behavior in the open field, but performed well in the active avoidance.

### 3.2. Augmented beta/gamma oscillations in the ACC of GAD65-/- mice

ACC oscillations were analyzed *ex vivo* through the activation of cholinergic and glutamatergic transmission upon co-perfusion of carbachol and kainate. Oscillations obtained in this reductionist slice model have been shown to be modulated by behavioral manipulations such as chronic stress (Ito et al., 2020) and provide robust approach to study beta (10-25 Hz) and slow gamma range (26-45 Hz) oscillations as functional read outs of interneuron activity (Steullet et al., 2014). A two-way ANOVA analysis revealed significant interaction of sex x genotype on the peak frequency at the beta range (Fig. 4D; two-way ANOVA for 10-25 Hz: interaction of sex x genotype: F(1,139)=5.088, p=0.0256), but a paired comparison for each sex did not demonstrate significant differences between female or male GAD65+/+ vs. GAD65-/- mice. At the slow gamma range at 26-45 Hz a significant effect of sex was observed with slightly increased peak frequencies for male compared to female mice (Fig. 4D; two-way ANOVA for 26-45 Hz: effect of sex: F(1,139)=4.016, p=0.047). The integrated power of oscillations at 10-25 Hz and 26-45 Hz was increased in GAD65-/- mice of both sexes (Fig. 4E; genotype effect for 10-25 Hz: F(1,139)=11.42, p=0.0009; 26-45 Hz: F(1,139)=4.594, p=0.034), without further effects of sex or interactions of sex x genotype.

**Figure 4:**
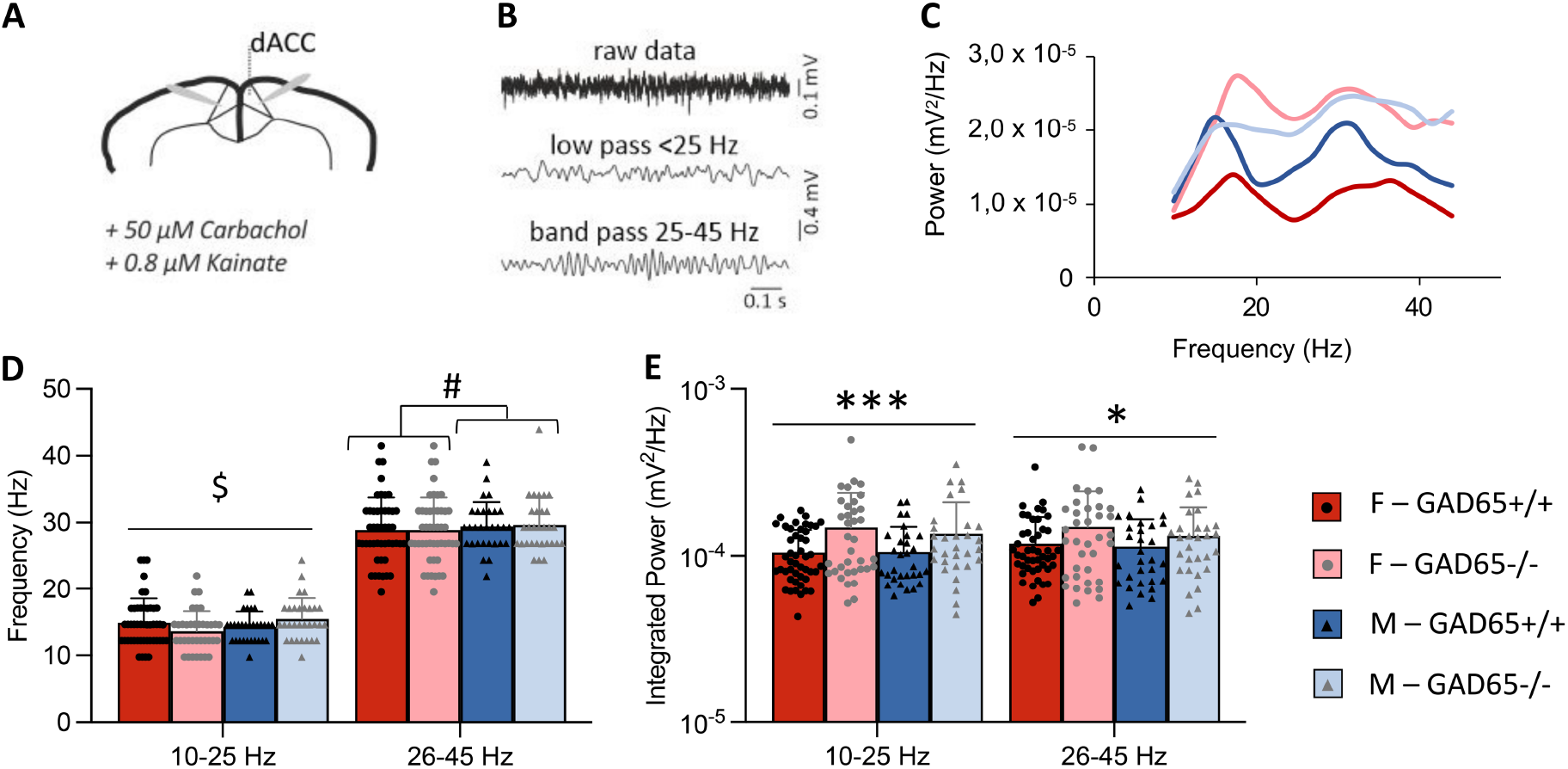
Increased propensity for induced fast oscillations in the ACC of GAD65-/- mice. **(A)** Scheme of an ACC slice and electrode positions for network oscillation recording induced by carbachol (50 µM) and kainate (0.8 µM) administration**. (B)** Representative traces of unprocessed data (5 kHz), low-pass filtered (<25 Hz) for illustration of beta-dominant LFP and band-pass filtered (25-45 Hz) for illustration of slow gamma oscillations *in vitro*. **(C)** Example power spectra of female and male mice across oscillation frequencies with two main peak frequencies in the LFP signal; approx. at 15 Hz and 35 Hz. **(D)** A significant interaction of sex x genotype was observed for the peak frequency at 10-25 Hz, but without any differences revealed by paired comparison. A sex effect was evident for the peak frequency at 26-45 Hz. **(E)** The integrated power of fast oscillations at the 10-25 Hz and 26-45 Hz frequency ranges were increased in GAD65-/- mice compared to their wildtype littermates, irrespective of their sex. All data presented as mean + SD with superimposed individual values. Two-way ANOVA: $ significant interaction of sex x genotype, p<0.05; # significant effect of sex, p<0.05; * significant effect of genotype, p<0.05; *** p<0.001.

*Ex vivo* gamma oscillations induced by CCh (5 µM) were further assessed in slices containing the ventral-to-intermediate CA3 and CA1 subregions (see supplementary material, Fig. S1). Here, the peak frequency of gamma oscillations (20-80 Hz) was significantly modulated by an interaction of sex x genotype in vCA3 (Fig. S1C; two-way-ANOVA: interaction of sex x genotype: F(1,70)=6.19, p=0.015), with a significant effect of genotype (F(1,70)=4.676, p=0.034), but not sex. Paired comparison revealed that male GAD65-/- mice showed higher gamma oscillation peak frequencies compared to male GAD65+/+ littermates (Fig. S1C; Student’s T-test: T(31)=3.274, p=0.003), while females showed comparable levels. The peak frequency in vCA1 was not significantly affected by sex, genotype nor by their interaction (Fig. S1C). A significant interaction of sex x genotype was observed for the integrated power of gamma oscillations in vCA3 as well (Fig. S1D; two-way-ANOVA: interaction of sex x genotype: F(1,70)=4.842, p=0.0311), but a paired comparison of genotype effects within each sex did not reveal significant differences. No significant effects were observed for the integrated power in vCA1 (Fig. S1D). Of note, the synchronization of gamma activity between vCA3 and vCA1 was significantly affected by sex (Fig. S1E; two-way-ANOVA, effect of sex: F(1,65)=9.598, p=0.003).

Together, electrophysiological recordings demonstrate a rather genotype-mediated effect for ACC beta/gamma oscillations (increased oscillation power in both male and female GAD65 -/- mice), while hippocampal gamma oscillation properties appear driven by sex differences (increased CA3-CA1 gamma correlation in males). These results suggest a potentially altered interneuron function in these brain regions relevant for emotional behaviors due to their involvement in generation of fast LFP oscillations.

### 3.3. Altered interneuron density in anxiety-related brain areas of GAD65-/- mice

Oscillatory activity is shaped by the activity of different interneuron subpopulations, mainly by PV-positive interneurons (Gulyás et al., 2010). Therefore, the number of these interneurons as well as SST-positive cells for comparison were analyzed in brain areas relevant for emotional behavior. Two-way ANOVA revealed an increased number of PV-positive cells in the ACC of GAD65-/- mice regardless of sex (Fig. 5A; effect of genotype: F(1,20)=8.301, p=0.009). The number of GAD67-postive cells, another isoform of GAD found ubiquitously in the cytosol of interneurons, was reduced in GAD65-/- mice (Fig. 5A; Mann-Whitney for main effects for non-normally distributed data: genotype: U=-2.57, p=0.008). No significant change in the number of SST-positive cells was found in this area (Fig. 5A).

**Figure 5:**
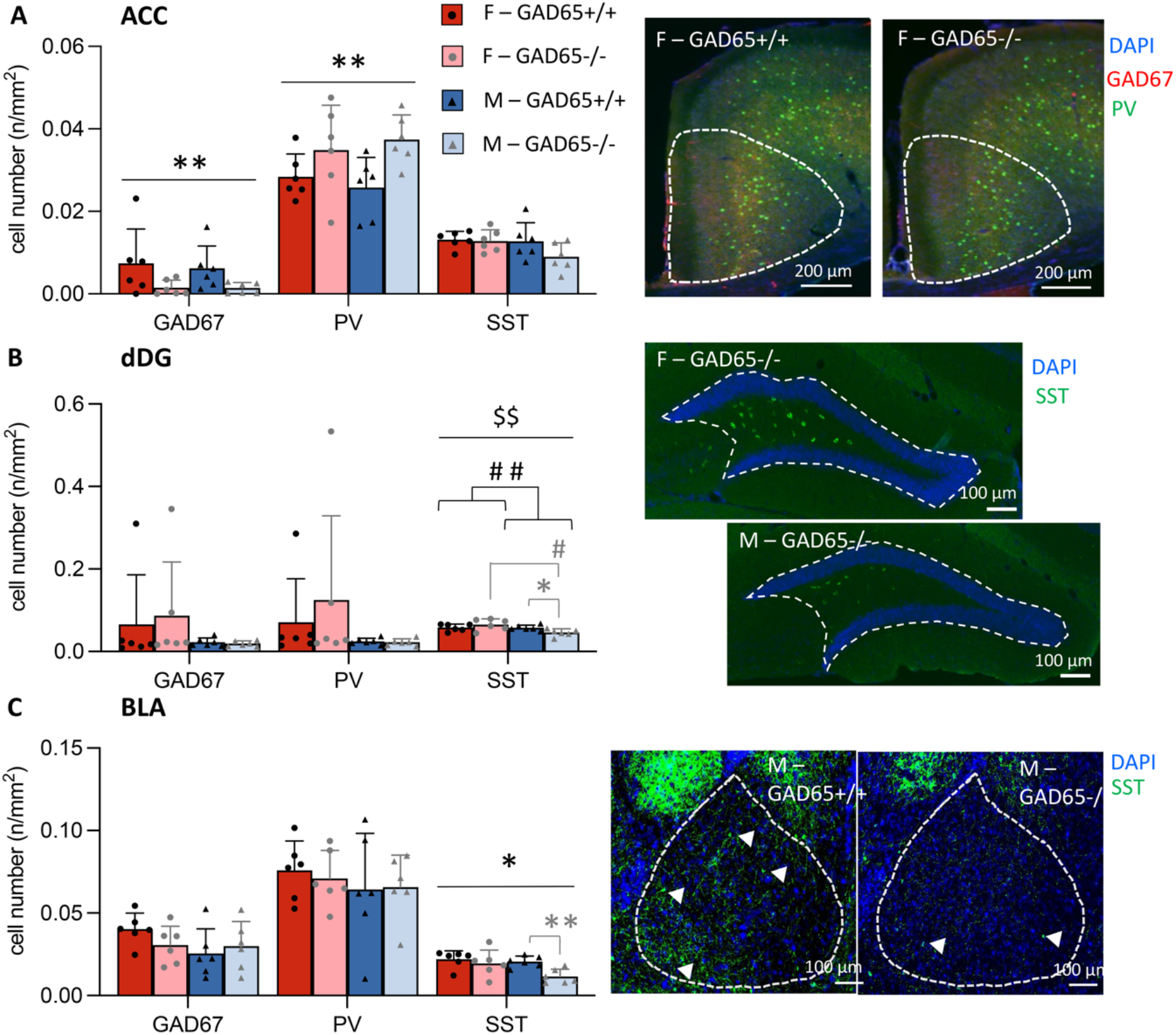
Altered interneuron density in anxiety-related brain areas of GAD65-/- mice. **(A)** In the posterior part of the anterior cingulate cortex (ACC), the number of GAD67- and parvalbumin (PV)-positive cells was increased in GAD65-/- mice of both sexes. Representative images of PV and GAD67 double immunostainings in the posterior ACC are shown on the right side. **(B)** In the dorsal dentate gyrus (dDG) the number of somatostatin (SST)-positive interneurons was generally lower in males, but interactions of genotype x sex were revealed. Paired comparisons demonstrated lower cell counts especially in male GAD65-/- mice. Representative images of SST immunostainings in the dDG are shown on the right side**. (C)** In the basolateral amygdala (BLA), SST-positive interneurons were generally reduced in GAD65-/- mice, but paired comparison confirmed that this cell population was reduced in especially in male GAD65-/- mice as well. All data presented as mean + SD with superimposed individual values. Two-way ANOVA: * significant effect of genotype, p<0.05, **p<0.01; # # significant effect of sex, ## p<0.01; $$ significant interaction of sex x genotype, p < 0.01. Paired comparison (T-Test): * (in grey) significant effect of genotype within the female subgroup, p<0.05, ** (in grey) p<0.01; # (in grey) significant effect of sex within the GAD65+/+ subgroup, p<0.05.

In the dDG, by contrast, a main effect of sex (Fig. 5B; F(1,20)=6.64, p=0.018) and a significant interaction of genotype x sex (F(1,20)=5.715, p=0.027) were observed for the number of SST-positive cells. Paired comparison revealed that male GAD65-/- mice had significantly less SST-positive cells in the DG compared to female GAD65-/- mice (T(10)=2.983, p=0.014) and to their male wildtype littermates (T(10)=2.633, p=0.025). The number of PV-positive and GAD67-postive cells did not differ in this area (Fig. 5B).

A main effect of genotype was observed for the number of SST-positive interneurons in the BLA (Fig. 5C; F(1,20)=6.486, p=0.019), while no significant effect of sex and no interactions of sex x genotype were found. However, as demonstrated by a paired comparison, a reduction of SST-positive cells was evident only in male GAD65-/- mice (T(10)=3.978, p=0.003) compared to male GAD65+/+ littermates. The number of PV-positive and GAD67-postive cells did not differ within the BLA (Fig. 5C).

In addition to interneuron counts, mRNA expression levels for PV and SST were assessed in the same areas in mice that went through the behavioral testing battery (see Supplementary material, Fig. S2). Here, no mRNA expression differences were observed in the ACC, but the expression of PV mRNA was reduced in the dDG of males (Fig. S2B; two-way ANOVA, main effect of sex: F(1,28)=6.973, p=0.013) as well as in the BLA of males (Fig. S2C; two-way ANOVA, main effect of sex: F(1,28)=5.112, p=0.036). In addition, expression of SST was increased in the BLA of male mice (Fig. S2C; Mann-Whitney for main effects for non-normally distributed data: main effect of sex U=2.569, p=0.009).

Together, the number of parvalbumin-positive cells was increased in the ACC of GAD65-/- mice regardless of sex, whereas the number of somatostatin-positive interneurons was reduced in the dDG and BLA of especially male GAD65-/- mice. No significant effects of genotype, sex or their interaction were observed on the number of PV- and SST-positive interneurons in vCA1 and vCA3 (Supplementary material, Fig. S3).

## 4. Discussion

In this study we investigated sex-specific differences in emotional information processing related to anxiety and threat avoidance in GAD65-/- mice that display a postnatal GABA deficit. To date, mainly male GAD65-/- mice have been investigated, showing increased anxiety, generalization of fear memory and deficits in fear extinction (see Müller et al., 2015 for review). However, epidemiological studies in human suggest sex differences with an increased incidence of anxiety disorders in women (McLean & Anderson, 2009; Vesga-Lopez et al., 2008). In accordance, in our rodent study, we found sex differences in anxiety, social preference and active avoidance learning that were partly modulated by GAD65 deficiency.

In an open field GAD65-/- mice of stayed more active over time, possibly indicating arousal or reduced habituation. This effect, indicated by an increased total distance covered, was more pronounced in females, adding to the previous observations of higher activity in male GAD65-/- mice compared to their wildtype littermates (Stork et al., 2000). Male GAD65-/- mice increased their activity in the center of the arena over time, suggesting an adaptive anxiety-like response. By contrast, with a short exposure and less aversive conditions under low light in the EPM, no difference in anxiety-like behavior was revealed and locomotor activity was even higher in females than males, regardless of the genotype. Previous studies indeed report that bright lighting conditions increase anxiety-like behavior and may even induce hyperactivity in males (Strekalova, 2005), in line with our observation in male wildtypes. Increased anxiety can potentially interact with cognitive tasks such as the novel object recognition test. Here, female wildtype mice showed no increased interest in a new object and male mice showed even a negative discrimination index that may indicate avoidance of novel objects. Thus, with the current setting of the NOR test, conclusions regarding recognition memory cannot be drawn. A careful adaptation of the protocol with an extensive pre-habituation to objects and test environments to overcome neophobia may allow assessing cognition with this test in future. When introducing social interaction partners instead of unanimated objects, mice of both sexes and genotypes preferred the partner mouse, despite reduced overall interaction times in female GAD65-/- mice (data not shown). In line with previous studies (Stack et al., 2010), male mice showed greater social preference that may also relate to aggression and an intensified interaction with the partner mouse restricted to a test tube. Social preference was enhanced in GAD65-/- mice compared to their wildtype littermates. In previous, studies male GAD65-/- mice showed reduced aggressive encounters that could be linked to increased anxiety as well (Stork et al., 2000), but in our setting with restricted access to interaction partners GAD65-/- mice may adapt their anxiety-like response. To assess threat-related behavior and learning in a more emotionally salient situation, a two-way avoidance paradigm in a shuttle box was used. Here, sex effects were revealed with higher levels of escapes in males over training session, in accordance with previous studies females performed better in the active avoidance task (Dalla and Shors, 2009). Variability was evident for avoidance responses, in accordance with previous observations for this task identifying faster and slower learning types (Galatzer-Levy et al., 2014). Based on such observations, fast learners were identified based on whether they showed more than 50% of successful avoidance trials (above chance level) already on the 2nd training session. 75% of female GAD65+/+ mice fell into this category, but only 25% of the male GAD65-/- mice, suggesting a discrete avoidance learning deficits in these animals. High avoidances on the 2^nd^ day as an approximation for fast learning did not correlate with fecal boli counts. Fecal boli can be assessed as a parameter for arousal (Hadad-Ophir et al., 2017; Sturman, 2018). Successful avoidance training may facilitate adaptation of arousal during this emotionally more demanding task by gaining cognitive control, resulting in reduced boli counts that occur after several days of training (Hadad-Ophir et al., 2017). Our results suggest that a high avoidance rate already in early training sessions does not predict an adaptation of arousal. Female GAD65+/+ mice had lower boli counts already at the beginning of the training, while female GAD65-/- mice adapted over time and male mice of both genotypes showed higher boli counts throughout the training sessions. To assess whether arousal or a priori differences in stress axis responsiveness may contribute to possible sex and genotype differences would be an interesting topic for future studies, for example by measuring cortisol and other stress-related hormones. Indeed, recent studies demonstrate that male GAD65-/- mice show higher corticosterone levels even under baseline conditions (Albrecht et al., 2022).

Adapting cognitive demands in emotional challenging situations relies on the function of a limbic network comprising the amygdala, the hippocampus and their interaction with frontal cortical areas. The ACC is an important part of the anxiety network and involved in the assessment of emotional situations and the appropriate choice of action, especially in stressful situations (Walton, 2007). It is active during threat perception (Straube, 2009) and a hyperactivity of this region, together with the amygdala and the ventral prefrontal cortex, is observed in patients with a generalized anxiety disorder (McClure et al., 2007). During attentional processes, oscillations at the gamma frequency range are registered in the frontal cortex and the hippocampus (Voloh et al., 2015; Rothé et al., 2011). To test whether oscillatory activity at gamma range is also altered in the ACC of male and female GAD65-/- and GAD65+/+ mice, we used a reductionist slice model that depends on activation of cholinergic and glutamatergic transmission upon co-perfusion of carbachol and kainate (Steullet et al., 2014). *Ex vivo* ACC oscillations are sensitive to behavioral manipulations such as chronic stress (Ito et al., 2020) and were robustly induced at the beta (10-25 Hz) and slow gamma range (26-45 Hz) in slices of all groups (Fig. 4). Phasic inhibition provided by PV-positive interneurons plays a fundamental role in sustaining and pacing *ex vivo* LFP oscillations in the hippocampus (Gulyas et al., 2010) but also in various cortical structures including ACC (Cabungcal et al., 2013; Steullet et al., 2014). Specifically, enhanced *ex vivo* beta/gamma power in the ACC was observed upon degradation of the perineuronal network that enwraps PV-positive interneurons (Steullet et al., 2014) and low PV-positive cell number was associated with reduction of *ex vivo* ACC beta/gamma oscillation power (Cabungcal et al., 2013). In accordance with these observations, we found an enhanced beta/gamma oscillation power and increased number of PV-positive interneurons in the ACC of GAD65-/- male and female mice. Interestingly, the observed reduction in the density of GAD67, an isoform producing GABA in the cytosol, in the ACC of GAD65-/- mice may suggest a generally increased excitability via disinhibition of excitatory pyramidal neurons which could also contribute the enhanced LFP oscillations in the ACC. It appears that GAD67 cannot fully compensate for GAD65 deficiency, resulting in the reduced GABA content in cortical areas of these mice (Stork et al, 2000). In the ventral hippocampus augmented gamma oscillations have been reported upon elevation of anxiety levels and aberrant gamma hypersynchrony is linked to symptom severity in post-traumatic stress disorder (PTSD) (Çalışkan and Stork, 2019; Dunkley et al., 2014). Effects of GAD65 deficiency in this network activity were minor, but we observed an enhanced CA3-CA1 gamma hypersynchrony in male compared to female mice of both genotypes. This sex effect may be linked to increased anxiety-like behavior of males in specific tests, such as decreased EPM activity and higher escape responses during avoidance learning. No sex-specific effects on the number of PV-positive cells were observed in the ventral CA3 or CA1 area, but the number of SST-positive was reduced in the dDG of males as well as in the BLA of especially in male GAD65-/- mice. The DG is relevant for active avoidance learning and anxiety-like behavior as well and a recent study using local knock down strategies support a role for GAD65 in these behaviors (Tripathi et al., 2021). Moreover, previous studies demonstrate anxiolytic actions of the peptide SST via the BLA (Gaskin et al., 2021) and analyzing mRNA levels four days after the last active avoidance training (Fig. S2) revealed a small increase in SST levels in the BLA of male mice, paralleled by decreased PV mRNA in this region.

Finally, we identified an increase in peak frequency of ACC (males and females) and hippocampal CA3 (only males) gamma oscillations in GAD65-/- mice which indicates faster gamma oscillation cycles via altered excitatory-inhibitory interactions. Here, a more detailed cell-type-specific analysis of GABA-A receptors subtypes that determine the frequency of fast oscillations (Heistek et al., 2010) might help to further resolve cellular correlates of gamma peak frequency alterations in the ACC and hippocampal CA3.

## 5. Conclusions and further perspectives

In summary, our analysis of behavior and network activity as well as interneuron density in male and female GAD65 mice revealed sex-dependent changes in activity, anxiety-like behavior and active avoidance learning that may rely on a network involving the BLA, dorsal and ventral hippocampal regions. This network may be further modulated by the ACC, which is driven by GAD65-dependent fast oscillatory activity (see Table 1 for summary of the results). Due to a possible stress induction by taking vaginal smear samples for estrus analysis in female mice, we refrained from correlating behavior, neuronal activity and expression values with estrous stages. Nevertheless, studies in cell culture and in vivo confirm that estrogen and its derivates can regulate the expression levels of GAD65 and other important co-neuromodulators in interneurons such as NPY or PV activity (Nakamura et al., 2004; Nakamura & McEwen, 2005; Ikeda et al., 2015; Sandhu et al., 2015; Picard et al., 2019). Therefore, we cannot rule out that individual variability in females in the analyzed responses might stem from different stages of the estrous cycle. Future studies are required that systematically address estrous phase effects on interneuron activity in limbic brain regions and their impact on anxiety-like and avoidance behavior. In addition, stress-related hormones such as corticosterone were not addressed in this study as well due to the invasive nature of repeated measurements, but they may provide interesting insights into the dynamics of stress responsiveness and behavioral adaptation. Moreover, this study engaged transgenic mice with a brain-wide GAD65 deficiency only on a C57Bl6/BomTac genetic background. Recent studies demonstrated that sex differences in behavioral traits such as activity, anxiety and social interaction vary across genetic backgrounds (Tabbaa et al., 2023), which should be considered as well in further experiments. Using local viral knock tools would allow to assess regional effects of an altered GAD65 expression in different mouse strains without time-consuming backcrossing breedings. Current analysis for such local intervention should be expanded to females (e.g. Tripathi et al., 2021; Albrecht et al., 2022).

Nevertheless, using a transgenic GAD65 deficient mouse allows to study genetic variation in GABA levels and their relevance for the function of anxiety networks in emotional challenging situations. In our animal model of lowered GABA levels due to GAD65 deficiency, we found that female GAD65-/- mice showed a more stable emotional response in challenging situations compared to males. These observations may relate to reduced active coping and increased anxiety-like behavior in male mice, while in humans, anxiety endophenotypes and anxiety disorders are more prevalent in women. Despite general concerns of translatability of animal models to human conditions (e.g. Barroca et al., 2022), our set of data illustrates the complex interactions of sex and the GABAergic system in modulating adaptive behaviors in challenging situations and might be useful for deciphering neuronal correlates. Adaptation to emotionally challenging situations in rodents and humans is highly dependent on genetic factors, previous experience as well as the exact nature of the challenge. In humans various SNPs that affect GABA levels are described. One of those variants results in increased GABA levels especially in the ACC, as revealed by fMRI-SPECT, and is associated with increased harm avoidance behavior in female but not male participants as an anxiety-related trait (Colic et al., 2018). In male participants higher GABA levels had only a limited impact on ACC activation and a defensive coping style. In our mouse model, we observe reduced GABA levels due to GAD65 deficiency, which appears to affect anxiety and threat avoidance behavior to a greater extend in male mice. Thus, in humans as well as in rodents, females may regulate their avoidance behavior differently from males in emotionally challenging situations, at least in part, via a GABAergic impact on their anxiety network structures in a potentially dose-dependent fashion. Interestingly in female, but not male healthy volunteers ACC volume and positive coping style predicted lower anxiety- or depression-related psychopathologies (Holz et al., 2016), pointing towards an important modulatory role of the ACC in behavioral adaptation, in line with our data. Further studies in healthy volunteers and anxiety patients as well as targeted intervention studies in relevant animal models are needed to investigate such sex-dependent biases in ACC function and its interaction with other structures such as the hippocampus and BLA and to investigate a possible modulation of such anxiety networks by GABAergic neurotransmission.

## Abbreviations

AA: active avoidance
ACC: anterior cingulate cortex
BLA: basolateral amygdala
DAPI: 4, 6-diamidino-2-phenylindole
dDG: dorsal dentate gyrus
EPM: elevated plus maze
F: female
LFP: local field potentials
GAD65: glutamic acid decarboxylating enzyme, 65 kDa isoform
GAD67: glutamic acid decarboxylating enzyme, 67 kDa isoform
M: male
NOR: novel object recognition
OF: open field
PV: parvalbumin
SI: social interaction
SST: somatostatin
vCA1+3: ventral cornu ammonis 1+3.

## Acknowledgement

We thank Franziska Blitz, Antje Koffi von Hoff, Vivian Sand and Annika Lenuweit for excellent technical assistance.

