## Supplementary Material for "Sex differences in anxiety and threat avoidance in GAD65 knock-out mice"

** corresponding author:* Anne Albrecht

### Supplementary Material

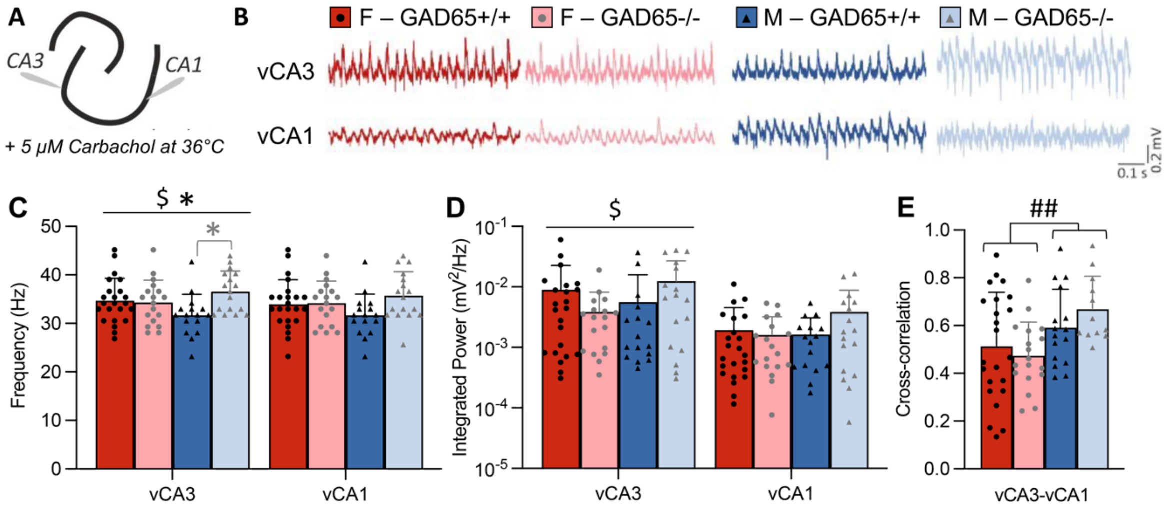


**Suppl. Fig. S1: Sex-specific alterations in gamma oscillations in the ventral CA3.** Gamma oscillations were induced in horizontal slices containing the ventral-to-mid hippocampal *Cornu ammonis* (CA)3 and CA1 regions by 5 µM Carbachol. **(A)** Scheme of a ventral hippocampal slice and electrode positions for network oscillation recording. **(B)** Example traces of carbachol-induced gamma oscillations in the CA3 and CA1 regions. **(C)** A significant interaction of sex x genotype and an effect of the genotype was observed for the frequency of ventral CA3 gamma oscillations. Paired comparison revealed a specific increase of the gamma frequency selectively in male GAD65-/- mice compared to male GAD65+/+ mice. **(D)** Within vCA3, a significant interaction of sex x genotype was found for the integrated gamma power (20-80 Hz), while neither frequency nor integrated power was significantly affected by GAD65 deficiency or by sex in the ventral CA1 region. Of note, the y axis is shown in logarithmic scale (log 10). **(E)** However, the gamma synchronization across CA3 and CA1 regions determined by cross-correlations was enhanced in male compared to female mice irrespective of the genotype. All data presented as mean + SD with superimposed individual values. Two-way ANOVA: $ significant interaction of sex x genotype, p<0.01; * significant effect of genotype, p<0.05; ## significant effect of sex, p<0.01. Paired comparison (T-Test): * (in grey) significant effect of genotype within the male subgroup, p<0.05.

*Methods in brief:* Horizontal slices containing the ventral to intermediate hippocampus were prepared from female GAD65+/+ (animals: N=5/ slices: n=23), female GAD65-/- (animals: N=5/ slices: n=18), male GAD65+/+ (animals: N=5/ slices: n=16) and male GAD65-/- mice (animals: N=4/ slices: n=17) as described previously (Çalışkan et al., 2016). Briefly, slices of 400 µm thickness were obtained with an angle of ~12° in the anterior-posterior direction to maintain the connectivity between the hippocampus and the entorhinal cortex and the CA3 and 1 subregions (Boulton et al., 1992; Mizunuma et al., 2014). The slices were then immediately transferred to an interface chamber perfused with carbogenated standard ACSF solution (rate ~2 ml/min, temperature 32 ± 1 °C). After at least 1.5 h of incubation, gamma oscillations were pharmacologically induced by adding 5 µM CCh for at least 45 min (Fisahn et al., 1998) and by increasing the temperature to 36 ± 1 °C. Recording electrodes were placed in stratum pyramidale of CA3 and CA1 regions in depth of ~80 µm and the network activity was recorded for at least 10 min. For statistical comparison, only slices showing signals with peak power higher than $10^{-5}$ $mV^2$ and peak frequencies higher than 20 Hz were included.

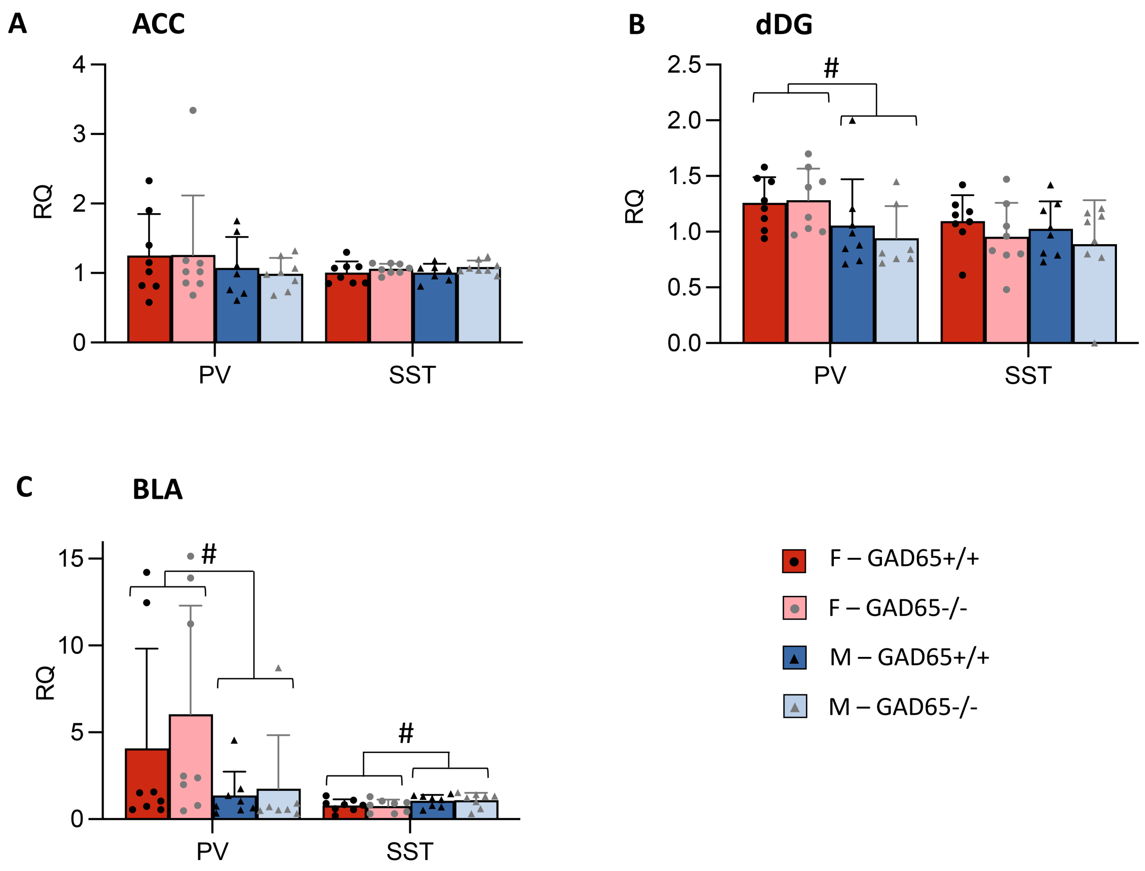


**Suppl. Fig. S2: Altered mRNA expression levels pf parvalbumin and somatostatin in anxiety-related brain regions after behavioral testing.**
The mRNA expression levels of parvalbumin (PV) and somatostatin (SST) were compared in a subset of mice that have been submitted to the behavioral test battery previously (n=7-8 per group). (A) No mRNA expression differences were observed in the ACC, but (B) the expression of PV mRNA was reduced in the dDG of males of both genotypes compared to their female counterparts. (C) A similar sex effect on PV expression was also observed in the BLA. In addition, expression of SST was increased in the BLA of male mice. Neither an effect of GAD65 deficiency nor a sex x genotype interaction was observed for mRNA expression levels.
All data presented as mean + SD with superimposed individual values. Two-way ANOVA: # significant effect of sex, $p<0.05$.

*Method in brief:* Animals were sacrificed four days after the last avoidance session by cervical dislocation under isoflurane anesthesia to analyse gene expression of somatostatin (SST) and parvalbumin (PV) in the posterior part of the anterior cingulate cortex (ACC, starting 1.10 mm from Bregma), the dorsal dentate gyrus (dDG) and basolateral amygdala (BLA, starting at -1.34 mm from Bregma, Paxinos and Franklin, 2001). Tissue was harvested with a micropuncher on coronary cut brains in a cryostat as described previously (Albrecht et al., 2022). Total RNA was isolated from tissue samples using the RNeasy Micro Plus kit (Qiagen, Hilden, Germany) according to manufacturer’s instructions. First-strand cDNA was synthesized with the Takara PrimeScript RT-PCR Kit (Takara Bio, Shiga, Japan), using 2.5 µM Oligo (dT) and 20 µM random hexamer first strand primers according to manufacturer’s instructions. Triplicates of a 1:5 dilution of cDNA were utilized for real-time PCR (ABI Prism Step One real-time PCR apparatus, Life Technologies) with TaqMan® reagents for predesigned assays of the target genes (assay

IDs: parvalbumin Mm00443100_m1; somatostatin Mm00436671_m1) and of the housekeeping gene Glycerinaldehyd-3-phosphat-Dehydrogenase (GAPDH; endogenous control; Life Technologies) in a duplex runs using two different fluorescent dyes (50 cycles of 15 s at 95°C and 1 min at 60°C, preceded by a 2 min 50°C decontamination step with Uracil-N-glycosidase and initial denaturation at 95°C for 10 min). Gene expression was normalized to male GAD65+/+ mice via the ddCT method by first normalizing the mean cycle threshold (CT) to the internal control GAPDH for each sample (dCT; dCT[target gene]) = (CT [target gene]) – (CT [GAPDH]) and then to the mean of the M – GAD65+/+ group with ddCT = dCT(sample) – mean dCT (M – GAD65+/+ group). Relative quantification (RQ) values were obtained by calculating RQ = (2-ddCT).

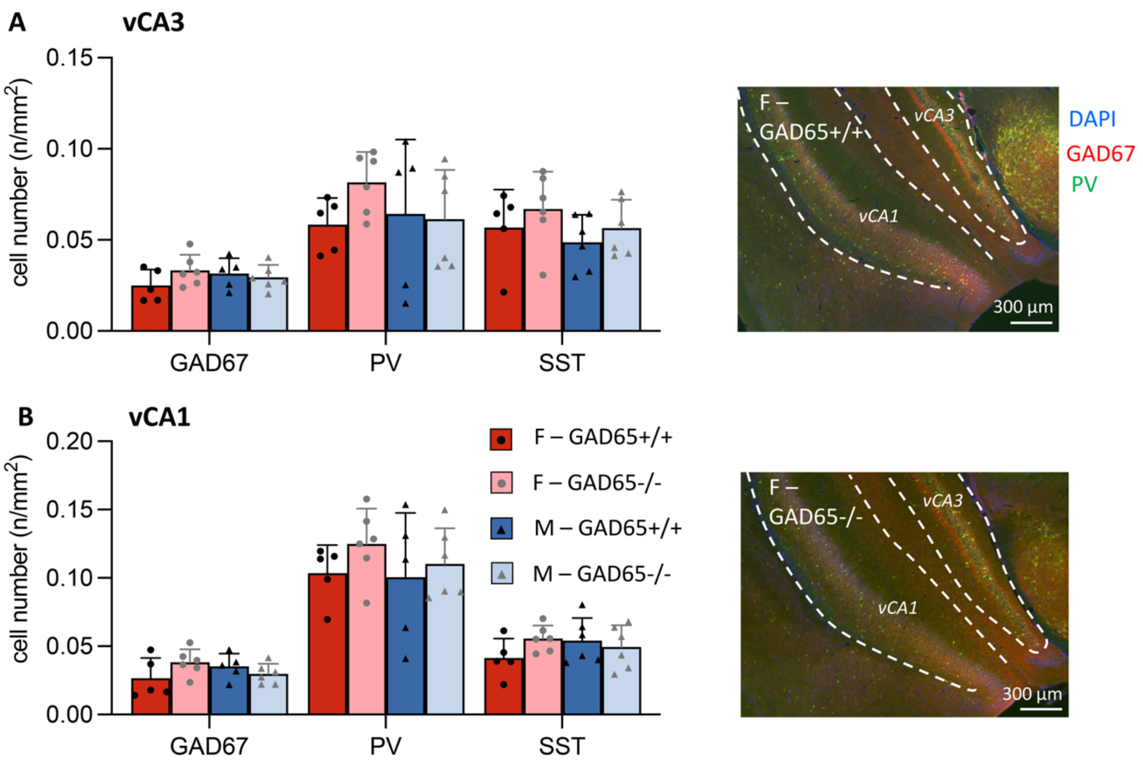


**Suppl. Fig. S3: No differences in interneuron density in the ventral hippocampus.**
Next to anterior cingulate cortex, dorsal dentate gyrus and basolateral amygdala (see Fig. 5), also coronal sections containing the ventral *Cornu Ammonis* region (vCA) 3 and 1 were immune-stained for GAD67 and Parvalbumin (PV) as well as for somatostatin (SST) and the number of positive cells was counted in the respective area. Example pictures for GAD67 and PV co-stainings are shown on the right. No significant effects of sex, genotype or their interaction were observed on cell counts in (A) the vCA3 or (B) the vCA1 region.
All data presented as mean ± SD with superimposed individual values.
